# Memory Transfer of Random Time Patterns Across Modalities

**DOI:** 10.1101/2020.11.24.395368

**Authors:** Kang HiJee, Auksztulewicz Ryszard, Chan Chi Hong, Cappotto Drew, Rajendran Vani Gurusamy, Schnupp Jan Wilbert Hendrik

**Affiliations:** Dept. of Neuroscience, City University of Hong Kong, Hong Kong S.A.R; Sensory and Motor Systems Research Group, Korea Brain Research Institute (KBRI), Daegu, South Korea; Neuroscience Dept, Max Planck Institute for Empirical Aesthetics, Frankfurt, Germany; Laboratoire des systèmes perceptifs, Département d’études cognitives, École normale supérieure, PSL University, CNRS, 75005 Paris, France

**Author notes:** Equal contribution (Kang HiJee, Auksztulewicz Ryszard). Corresponding author: Jan W. H. Schnupp, Dept. of Neuroscience, City University of Hong Kong, 31 To Yuen Street, Kowloon, Hong Kong.

**Keywords:** implicit learning, memory transfer, electrocephalography, crossmodal mechanisms, temporal processing

## Abstract

Perception is sensitive to statistical regularities in the environment, including temporal characteristics of sensory inputs. Interestingly, temporal patterns implicitly learned within one modality can also be recognised in another modality. However, it is unclear how cross-modal learning transfer affects neural responses to sensory stimuli. Here, we recorded neural activity of human volunteers (N=24, 12 females, 12 males) using electroencephalography (EEG), while participants were exposed to brief sequences of randomly-timed auditory or visual pulses. Some trials consisted of a repetition of the temporal pattern within the sequence, and subjects were tasked with detecting these trials. Unknown to the participants, some trials reappeared throughout the experiment, enabling implicit learning. Replicating previous behavioural findings, we showed that participants benefit from temporal information learned in audition, and that they can apply this information to stimuli presented in vision. Such memory transfer was not observed from vision to audition. However, using an analysis of EEG response learning curves, we showed that learning temporal structures both within and across modalities modulates single-trial EEG response amplitudes in both conditions (audition to vision and vision to audition). Interestingly, the neural correlates of temporal learning within modalities relied on modality-specific brain regions, while learning transfer affected activity in frontal regions, suggesting distinct mechanisms. The cross-modal effect could be linked to frontal beta-band activity. The neural effects of learning transfer were similar both when temporal information learned in audition was transferred to visual stimuli and vice versa. Thus, both modality-specific mechanisms for learning of temporal information, and general mechanisms which mediate learning transfer across modalities, have distinct physiological signatures that are observable in the EEG.

## 1. Introduction

Sensory signals in each modality contain complex feature-based information. Unlike modality-specific features - such as sound frequency or colour spectrum - timing information is universal across modalities, given their temporal nature (Hardy and Buonomano, 2016; Lim et al., 2016). For efficient sensory perception, extracting statistical regularities based on stimulus features is essential (Sohoglu and Chait, 2016; Nobre and van Ede, 2018). However, it is unclear whether this ability is mediated by global mechanisms or differs across sensory modalities. For instance, absolute timing acuity greatly differs between modalities, with the auditory modality being more sensitive to timing information (Goodfellow, 1934; Grondin and Rousseau, 1991; Grondin, 1993; Grondin and McAuley, 2009). Previous human psychophysics studies have reported auditory dominance for temporal processing, especially when the information is presented in multiple modalities concurrently (e.g., Repp and Penel, 2002; Recanzone, 2003; Grondin and McAuley, 2009). Later neuroimaging studies reported involvement of auditory regions during visual timing processing, further implying the dominance of the auditory system for processing timing regardless of stimulus modality (Grahn et al., 2011; Kanai et al., 2011). However, other studies claim that the reports of auditory dominance could be driven by the experimental settings requiring timing sensitivity as a crucial factor (Grahn, 2012; Rammsayer, 2014).

Reflecting discrepancies in the empirical studies, conflicting theoretical models posit that temporal processing requires either dedicated (modality-specific) or generalised (cross-modal) mechanisms (Buonomano, 2000; Karmarkar and Buonomano, 2003; Ivry and Schlerf, 2008). It is widely claimed that sub-second interval encoding is processed in modality-specific regions where temporal patterns are spatially coded by dynamic changes of neural states (Buonomano and Maass, 2009; Hardy and Buonomano, 2016). An opposing view suggests that temporal information is processed by an internal “pacemaker” emitting timing information to memory, particularly for longer time intervals (scalar expectancy theory; Gibbon, 1977; Gibbon et al., 1984; Rammsayer and Ulrich, 2001; Karmarkar and Buonomano, 2007). While most previous work focuses on processing single time intervals or rhythmic time patterns associated with motor tasks, sensory inputs from natural scenes are often more complex, containing temporal structure in both sub-second and supra-second ranges. It is currently unknown whether temporal patterns in the supra-second range, composed of irregular sub-second range intervals, are treated independently across modalities or mediated by generalised mechanisms.

One recent human psychophysics study attempted to investigate implicit learning of timing information across modalities, covering audition, touch, and vision (Kang et al., 2018). Participants showed an improvement on a given task only for a specific random time pattern that re-appeared over trials, even in an unsupervised learning setting. These improvements were observed for all three tested modalities, suggesting rapid and implicit learning of arbitrary temporal structure. Regardless of better overall task performance in audition than vision, the learning effect was qualitatively similar. Interestingly, participants showed successful memory transfer from audition to other modalities, suggesting that the memory encoding of random time patterns may be mediated by crosstalk between modalities. However, it is unclear which brain regions and neural activity patterns are involved in memory transfer across modalities.

Here, we measured brain responses of human volunteers using electroencephalography (EEG), during a repetition detection task (Agus et al., 2010; Kang et al., 2018). Across experimental sessions, they were exposed to auditory or visual stimulus sequences with irregularly timed stimuli. Unknown to the participants, some temporal patterns repeated throughout the experiment, and we examined whether participants benefited from temporal information learned in one modality when they perform the same task in another. Here, we developed a novel method for analysing learning curves in sensor and source space, to investigate which brain regions mediate learning transfer across modalities. While our behavioural finding showed the transfer of temporal pattern learning only from audition to vision, EEG findings confirmed the transfer of temporal pattern learning from audition to vision as well as from vision to audition. The neural effects were reflected in beta frequency band modulation of activity in the right inferior frontal gyrus (rIFG), suggesting modality-general encoding of the temporal patterns transferred across modalities.

## 2. Materials and methods

### 2.1 Participants

A total of 24 participants enrolled in the study (12 females, 12 males; median age = 20 years, SD = 3.24 years). All participants self-reported as normal hearing and having normal or corrected-to-normal vision, with 3 self-reported as left-handed and the remaining as right-handed. Participants were pseudo-randomly assigned to one of two test groups (audition to vision vs. vision to audition) to maintain the gender ratio.

Participants gave informed consent to taking part in the experiments and received cash reimbursement for their time after participating in the study. All study protocols were approved by the Human Subjects Ethics Sub-Committee of the City University of Hong Kong.

### 2.2 Experimental design and statistical analyses

#### 2.2.1 Stimuli

Auditory or visual stimulus “sequences” (brief trains of clicks or visual flashes) were presented in training or testing blocks which are described in detail below. Each sequence consisted of a set of 0.2 ms rectangular stimulus pulses delivered at an average rate of 7 Hz. To generate pulse trains following a Poisson distribution (with a 10 ms refractory period), inter-pulse time intervals were drawn from an exponential distribution, and intervals less than 10 ms were discarded. The length of each sequence was limited to 2.4 s. Sequences either had random time intervals for the full duration (random patterns, P), or a half length (1.2 sec.) of random intervals seamlessly presented twice (repeated patterns, RP). While for most trials, both P and RP stimuli were generated afresh each time and occurred only once, one specific RP sequence (reference repeated patterns, RefRP) randomly re-occurred several times within the test block. As in Agus et al. (2010), the rationale is that repeats in RefRP trials should become easier to detect, even though the subjects are generally unaware that they had encountered the RefRP pattern before in the same block.

Test blocks were divided into two conditions: in the Transfer condition, RefRPs had an identical temporal structure for both modalities, while in the Control condition, the RefRP temporal structure differed between modalities (Fig. 1).

**Figure 1.**
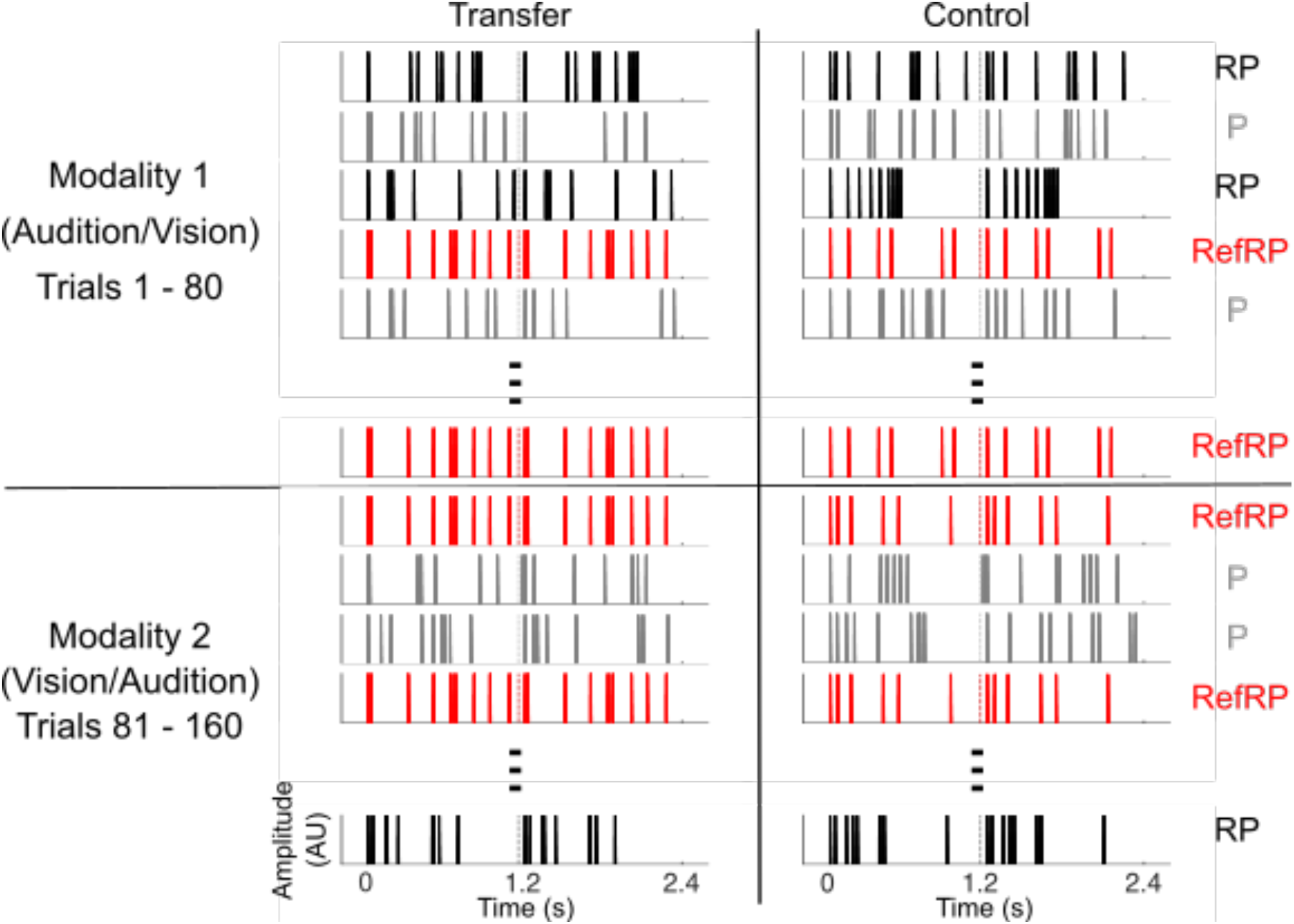
Example schematic of the stimulus patterns. Each stimulus example refers to one trial of either RP (black), P (grey), or RefRP (red). Different trial types were presented in a randomized order. Dotted lines within each stimulus indicate the midpoint of the pattern where the repetition begins for RP and RefRP trials. While stimuli are newly generated for RP and P, the RefRP is identical across trials in a block. The black solid horizontal line indicates the middle of the test block, at which point in time the stimuli are presented with a new modality. In the Transfer condition, the same RefRP is presented between two modalities, while in the Control condition, a different RefRP is presented after the modality switch.

In the auditory modality (Audition), the click trains were high-pass filtered at 2 kHz to avoid the possibility of spectral cues arising from filter ringing at a lower frequency range (Pollack, 1968; Kang et al., 2017). The low frequency range below 2 kHz was filled with band-passed pink noise between 50 Hz and 2 kHz at −2 dB SNR relative to pulse trains. The sound level for the auditory stimulus sequences was kept at 72 dB SPL, and they were presented at a 44.1 kHz sampling rate. In the visual modality (Vision) no filtering was applied to pulse trains, and identical filtered pink noise was presented acoustically during the visual stimuli via a separate channel for consistency between modalities. All stimulus sequence waveforms were generated using custom Matlab scripts (R2017a, MathWorks Inc.), and played via MOTU UltraLite-mk3 Hybrid soundcard. Auditory stimuli were delivered to the participants through Brainwavz B100 earphones.

For visual stimuli, a single green LED light was housed in a custom-made 7×4×3 cm box covered with translucent plastic film to diffuse dimmed light. It was attached on top of the screen at the subjects’ eye level. The LED lights were driven by a RM2 mobile processor (Tucker-Davis Technologies, TDT) and triggered by the MOTU soundcard.

#### 2.2.2 Apparatus and Procedure

Participants were seated in a sound-attenuated and electrically shielded room in front of a computer screen. The screen was used to deliver instructions and display a white fixation cross on a gray background during the auditory task. During the visual task, the screen was black and participants were instead asked to fixate on the LED delivering the visual pulse trains. After a given trial (auditory or visual), participants were asked whether the trial contained repetition within the sequence. Their responses were acquired by a keyboard button press on a testing laptop (1 for a repetition; 2 for no repetition). Before performing the actual task, all subjects received a short training session to confirm that they correctly understood the task. There were three training blocks for each modality. The first training block for each modality had 10 trials of either RPs with 10 repetitions of a short 1.2 sec. segment of pulse trains, or the equivalent length of a random non-repeating pattern (P). Under these conditions, the repetition in RP trials was very salient so that participants could familiarise themselves with the task. The second and third training blocks consisted of 20 trials each, with the number of repetitions within each RP trial first reduced to 3, and then to 2. During the training sessions, participants received feedback after each trial on whether they made a correct response. Since the training session was solely designed to ensure that participants understood the task requirements, no RefRP stimuli were included in the training. The training was done first on auditory, then on visual pulse trains. All subjects showed at least 50% correct responses in the final training block for both Audition and Vision. Average correct responses of the final training block were 69.8% for Audition and 66.2% for Vision.

Prior to performing the actual task, the participants were fitted with an EEG cap, and a conducting gel was applied on the scalp under each of the EEG electrode contacts. Participants were instructed to minimise blinking and body movement during stimulus presentation. EEG signals were recorded using ANT Neuro EEGo Sports 64 channel device at a sampling rate of 1000 Hz.

Participants were randomly assigned to one of two groups. One group performed the task in Audition first and then moved on to Vision (AV). The other group performed the tasks in the opposite order (VA). Each participant completed a total of 4 test blocks. Each block contained 160 trials −80 trials for the first modality and 80 trials for the second modality. Half way through each block, participants were instructed about the modality switch. Each modality contained 40 P, 20 RP and 20 RefRP trials, presented in a random order. Participants performed 2 blocks of each condition, Transfer and Control, in a randomised order. Each block contained a newly generated RefRP.

Two seconds after the onset of the stimulus sequence in each trial, a visual prompt emerged on the screen asking the subject to indicate whether they had perceived a within-sequence repetition after the sequence ended. No trial-by-trial feedback was provided during the actual task.

#### 2.2.3 Behavioural data analysis

First, based on signal detection theory, behavioural *d*’ sensitivity index was calculated per participant, separately for RefRP and RP stimuli which were generated in the same way but differed in its re-occurrences across trials. *d*’ values were then rescaled to the across-subject mean performance in each condition and modality order group relative to each subject’s overall mean performance, to minimise the individual performance difference and focus on the performance gain that could have occurred due to the learning of RefRP. To test whether participants were more sensitive to make correct responses for RefRP than RP, a mixed-design analysis of variance (ANOVA) was performed on *d*’ values using, with stimulus type (RefRP or RP), condition (Transfer or Control), and modality (Audition or Vision) as within-subject factors, order (AV or VA) as a between-subject factor, and participant as a random factor. Post-hoc repeated-measures ANOVAs were conducted separately for each level of the modality and order factors. Lastly, paired *t*-tests were conducted on the performance difference between RefRP and RP per modality and condition to see whether there was a behavioural benefit of receiving the same RefRP in the Transfer condition compared to the Control condition.

#### 2.2.4 EEG data analysis

The recorded EEG multichannel signals were preprocessed using the SPM12 Toolbox (Wellcome Trust Centre for Neuroimaging, University College London) for Matlab. The EEG data were high-pass filtered at a cut-off frequency of 0.01 Hz using a 5th order two-pass Butterworth filter to reduce slow drifts such as changes in impedance due to sweating. A 5th order two-pass Butterworth notch filter (48-52 Hz) was applied to remove line artefacts. Then, a 5th order two-pass Butterworth low-pass filter at 90 Hz was applied to reduce any high-frequency environmental or physiological noise. Eyeblink artefacts were removed by subtracting the first two spatiotemporal components associated with each eyeblink (Ille et al., 2002), as implemented in SPM12. Specifically, the first two principal components were extracted from the time course and topography of the average eyeblink-evoked response, and removed from the raw EEG data at the time of each blink using spatial filtering. After this preprocessing, the data were further denoised based on Dynamic Separation of Sources (de Cheveigné and Simon, 2008). This denoising procedure is commonly used to increase the signal-to-noise ratio of event-related potentials (ERPs) by maximizing the reproducibility of stimulus-evoked responses across trials. To calculate ERPs, continuous data were epoched between 200 ms before stimulus onset to 2600 ms after stimulus offset. Each epoch (segment) was baseline-corrected to the mean of the prestimulus period (i.e., from −200 ms to 0 ms relative to stimulus onset). In order to exclude epochs contaminated by transient artefacts, we removed epochs that had an average RMS (root mean square) amplitude exceeding the median by 2 standard deviations from further analysis.

The remaining epochs were subjected to single-trial learning-curve fitting. First, to determine the learning curve equation, we fitted four models to the grand-average ERP RMS amplitude. To calculate the grand-average data, first - per participant, session, stimulus type, and channel - a “relative RMS response amplitude” was calculated for each of the 20 trials of each stimulus type as RMS(stimulus)-RMS(baseline). Then, these single-trial relative RMS values were averaged across participants, sessions, stimulus types, and channels. The four models used to fit the grand-average data were based on previous studies in which learning curves were quantified in the context of statistical learning (Kepinska et al., 2017; Siegelman et al., 2018; Choi et al., 2020) and sensory adaptation (Ulanovsky, 2004). They included linear, quadratic, exponential, and logarithmic models, described by the following four equations respectively:

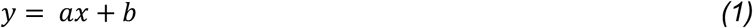

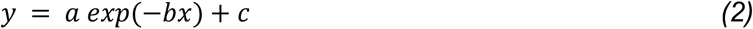

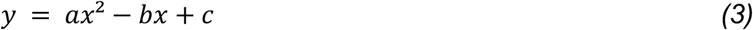

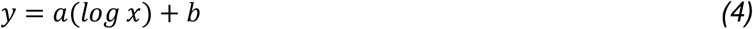

In equations (1)-(4), *x* is the trial index number (ranging from 1 to 20); *a*, *b* and *c* are model coefficients; and *y* is the grand-averaged RMS amplitude of all data at trial *x*. To select the most appropriate among these four candidate models, the models were compared using the adjusted R^2^ metric, which quantifies the goodness of fit of each model, while penalising for the number of model coefficients (Miles, 2014). The adjusted R^2^ values, grand-averaged across participants for each model, were: linear (eqn 1): 0.309; quadratic (eqn 2): 0.348; exponential (eqn 3): 0.340; and logarithmic (eqn 4): 0.384. Since the logarithmic model yielded the highest adjusted R^2^ on average, and to ensure that the learning curve coefficients are comparable across participants, it was selected for subsequent learning curve estimation in each individual participant and condition (Fig. 4A).

To estimate learning curves, epoched EEG data were sorted into 6 trial types per session according to two within-subjects factors - modality (Audition; Vision) and stimulus type (P; RP; RefRP). Additionally, across sessions, data were grouped according to a third within-subjects factor (Transfer; Control). Finally, across participants, data were grouped into a between-subjects factor of modality order (AV; VA). This yielded a 2×3×2×2 design with three within-subjects factors (modality, stimulus, condition) and one between-subjects factor (order). For each participant, channel, time point, and stimulus/trial type (design cell), singletrial EEG amplitudes corresponding to the first 20 trials of a particular stimulus/trial type were fitted with the logarithmic fit, yielding the logarithmic coefficient *a* (quantifying the “learning rate”) and a constant *b* (quantifying the “baseline” ERP amplitude, after regressing out the learning curve).

The resulting ERPs and learning curves were analysed separately, in a series of repeated-measures ANOVAs. Specifically, we designed separate flexible factorial designs (repeated-measures ANOVAs) for the following factors: dependent variable (ERP; learning curve); order (AV; VA), and modality (Audition; Vision). Each ANOVA had the within-subjects factors stimulus (P; RP; RefRP) and condition (Transfer; Control), as well as a random participant factor. ERP and learning curve data were converted into 3D images (2D: scalp topography; 1D: time), smoothed with a 8 mm x 8 mm x 50 ms Gaussian kernel (to ensure that data conform to random field theory assumptions, used in SPM12 to correct for multiple comparisons across time points and channels), and entered into the flexible factorial designs. Statistical parametric maps (SPMs) were thresholded at *p* < 0.005 (uncorrected) and significant effects were inferred at a cluster-level *p* < 0.05 family-wise error (FWE), correcting for multiple comparisons across time points and channels under random field theory assumptions (Kilner et al., 2005).

For each ANOVA, the following contrasts were examined: First, we looked at the main effects of stimulus (RefRP vs. RP: testing for the effect of stimulus repetition across trials). We focused on this comparison only, since the difference between RefRP and P stimuli includes multiple factors (repetition within trials, repetition between trials, as well as differences in button press preparation), while the difference between RefRP and RP only includes repetition between trials - i.e., our effect of interest in the context of pattern learning. Second, we examined the main effect of condition (Transfer vs. Control: a sanity check, testing for overall differences between experimental sessions). Finally, we tested the interaction effect of stimulus (RefRP vs. RP) and condition (Transfer vs. Control), zooming into the cross-modal learning transfer effects. We reasoned that, in the second modality, RefRP stimuli should be processed differently between Transfer and Control conditions, since in the Transfer condition the subjects had just experienced the same temporal pattern in a different modality, while in the Control condition they had not. Conversely, RP stimuli should be processed identically in the Transfer and Control conditions.

The significant interaction effects were subjected to an additional post-hoc analysis, verifying whether the effects of “learning” a RefRP pattern in the previous modality significantly affect the initial presentations of RefRP stimuli in the Transfer condition, in the second modality. This would indicate that the temporal pattern identifying the particular RefRP that had been learned in one modality could be applied in another modality. To this end, we investigated the differences in trial-by-trial changes in EEG signals between four trial types (stimulus: RefRP vs. RP; condition: Transfer vs. Control). Per participant and trial, we extracted data from EEG channels corresponding to the significant SPM cluster (using the 2D coordinates of each EEG channel mapped onto the 2D cluster coordinates), calculated the RMS of the significant time window (averaged across channels), and subtracted the RMS of the pre-stimulus baseline. Based on previous studies (Kang et al., 2017; 2018), we expected rapid learning to occur within at most the first five trials. To test this hypothesis, the RMS data were averaged across the first three trials of each trial type, normalised per participant by dividing each data point by the standard deviation (pooling across trial types), and subject to pairwise comparisons (paired *t*-tests).

The spatiotemporal clusters of significant effects were source-localised using the multiple-sparse-priors approach under group constraints (Litvak and Friston, 2008). For each significant main effect, sensor-level data were subject to source localisation for a time window in which the significant effect was observed, plus an additional 100 ms of signal on either side, given that, in multiple-sparse-priors source reconstruction, rise and fall times of neural signals should be included, as source activity estimates are based on signal variance across time (López et al., 2014). For each participant, the source estimates were converted into 3D images (in MNI space), smoothed with a 5×5×5 mm Gaussian kernel, and entered into flexible factorial designs with one within-subjects factor (Stimulus) and one between-subjects factor (Participant). The resulting statistical parametric maps were thresholded at *p* < 0.05 (two-tailed, uncorrected) and significant effects were inferred at a cluster-level *p* < 0.05 family-wise error (FWE, small-volume corrected), correcting for multiple comparisons across voxels under random field theory assumptions (Kilner et al., 2005). Sources were labelled using the Neuromorphometrics atlas, as implemented in SPM12.

In case of significant interaction effects, the source reconstruction procedure was identical as for the main effects, except that sensor-level data were used to calculate contrast time-series (RefRP-RP). Flexible factorial designs had one within-subjects factor (condition: Transfer vs. Control) and two between-subjects factors (participant; order: AV vs. VA). Statistical thresholds were set at *p* < 0.05 (uncorrected), correcting for multiple comparisons across time points at a cluster-level *p* < 0.05 family-wise error (FWE).

Furthermore, we investigated whether pattern learning transfer was reflected in changes in activity in specific EEG frequency bands. Since we observed significant interaction effects between stimulus and condition for both AV and VA participant groups, which in both cases source-localised to the right inferior frontal gyrus (rIFG; see Results), we focused on the time-frequency activity localized to the rIFG. To this end, continuous EEG data were re-epoched ranging from 800 ms before stimulus onset to 800 ms after stimulus offset. Dipole waveforms were estimated for rIFG coordinates (dipole location: peak MNI coordinates of the source reconstruction in each AV and VA group; dipole orientations: three orthogonal dipole moments), resulting in three dipole time-series (corresponding to three orthogonal dipole orientations) per trial. The dipole time-series were subject to a timefrequency estimation using multi-taper convolution with a frequency range of 8-48 Hz (frequency step: 1 Hz) and a time window of 400 ms (time step: 50 ms), yielding three timefrequency maps per trial. For each trial and time-frequency point, maximum power was selected across the three orthogonal dipole orientations, and log-rescaled to the pre-stimulus (600-200 ms) baseline. These single-trial rescaled time-frequency estimates were subject to logarithmic fitting, as described above.

The resulting time-frequency maps of model coefficients (learning curves) were converted into 2D (time x frequency) images per trial type, smoothed with a 5 Hz x 50 ms Gaussian kernel (to ensure that data conform to random field theory assumptions), and entered into a factorial design with two within-subjects factors (stimulus: RefRP vs. RP; condition: Transfer vs. Control) and one between-subjects factor (participant). Statistical parametric maps were thresholded at p < 0.05 (two-tailed, uncorrected) and significant effects were inferred at p < 0.05 family-wise error (FWE), correcting for multiple comparisons across time and frequency under random field theory assumptions (Kilner et al., 2005). As in the case of EEG amplitudes, the significant interaction effects were subject to an additional post-hoc analysis to test if the interaction between stimulus and condition modulates stimulus processing for the initial trials. Per participant and trial, we pooled power estimates from the significant cluster, averaged them across the first three trials of each trial type, and normalised the resulting estimates of initial power per participant by dividing each data point by the standard deviation (pooling across trial types). These data were subject to pairwise comparisons (paired *t*-tests).

Finally, we tested whether behavioural sensitivity correlates with neural signatures of learning transfer (i.e., the effects of learning transfer on EEG amplitude and time-frequency activity) across participants. As the behavioural measure, per participant and condition (Transfer vs. Control) we calculated the difference in d’ between RefRP and RP stimuli presented in the second modality. As the neural signature of learning transfer on EEG amplitude, we calculated the differences in EEG-based logarithmic fits (pooling from significant clusters shown in Fig. 4BC and 4EF; see Results) between RefRP and RP stimuli, separately for each participant and condition (Transfer vs. Control). Similarly, as the neural signature of learning transfer on time-frequency activity, we calculated the corresponding differences in beta-band logarithmic fits (pooling from significant clusters shown in Fig. 5B). The single-trial estimates of differences in behavioural sensitivity to RefRP and RP stimuli were entered into an analysis of covariance (ANCOVA) with the following independent variables: (1) EEG signature of learning transfer (as a continuous regressor), (2) timefrequency signature of learning transfer (as a continuous regressor), (3) condition (Transfer vs. Control), (4) order (AV vs. VA), as well as their interactions. Significant interactions were inspected using post-hoc ANCOVAs.

**Figure 2.**
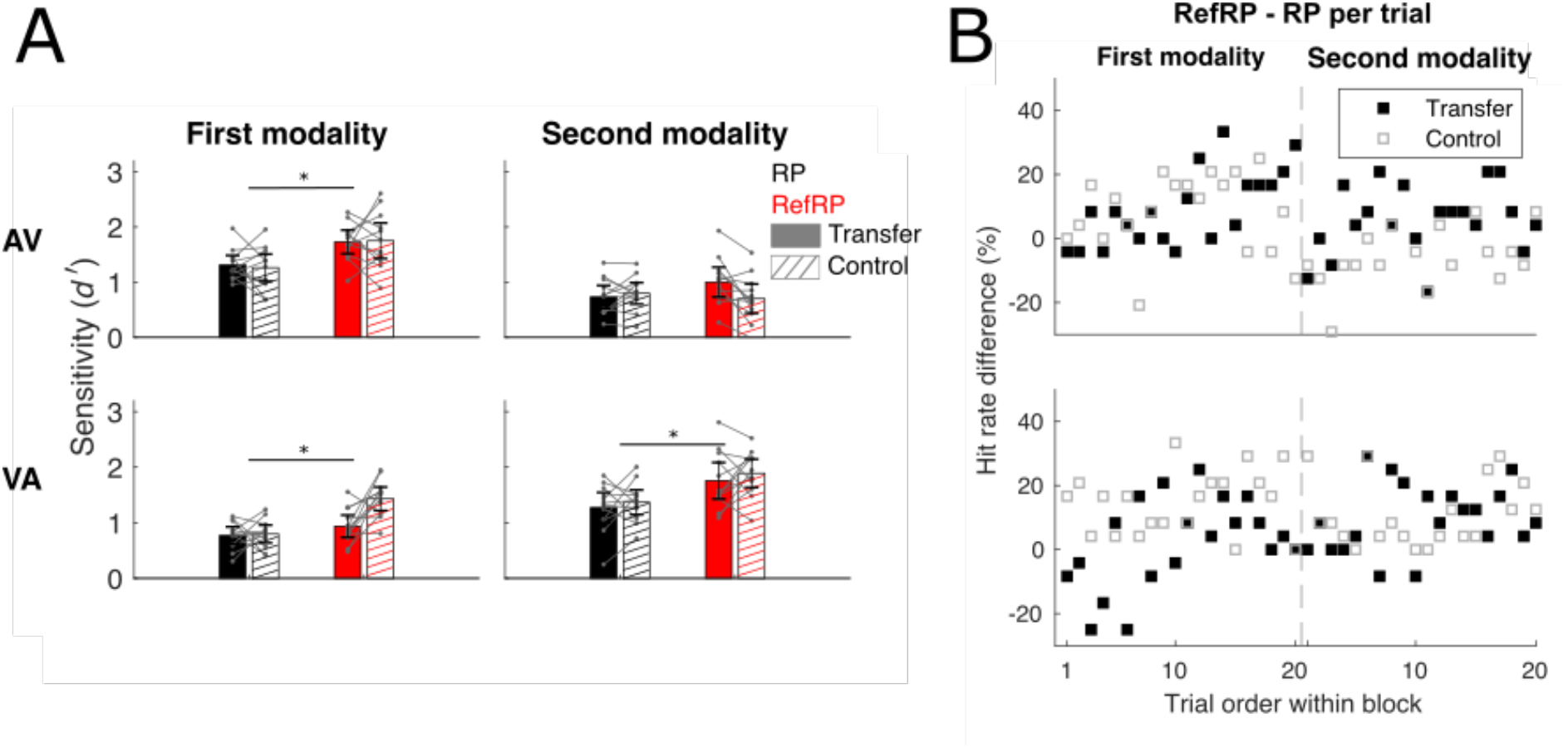
Behavioral results. **(A)** Sensitivity index *d*’ values averaged across subjects for RP (black) and RefRP (red) for each modality (top: AV group, bottom: VA group) and each condition (filled: Transfer, hatched: Control). Grey lines on the bar graphs indicate individual subject’s *d*’ values per stimulus type and condition. Error bars represent 95% confidence intervals. Asterisks indicate where there are statistically significant differences between stimulus types (*p* < 0.05 to indicate main effect between stimulus types). **(B)** The time course of performance difference (RefRP - RP) for Transfer (filled markers) and Control (empty markers) conditions (top: AV group, bottom: VA group). Dashed lines indicate the midpoint of the test block where the modality switch occurred.

**Figure 3.**
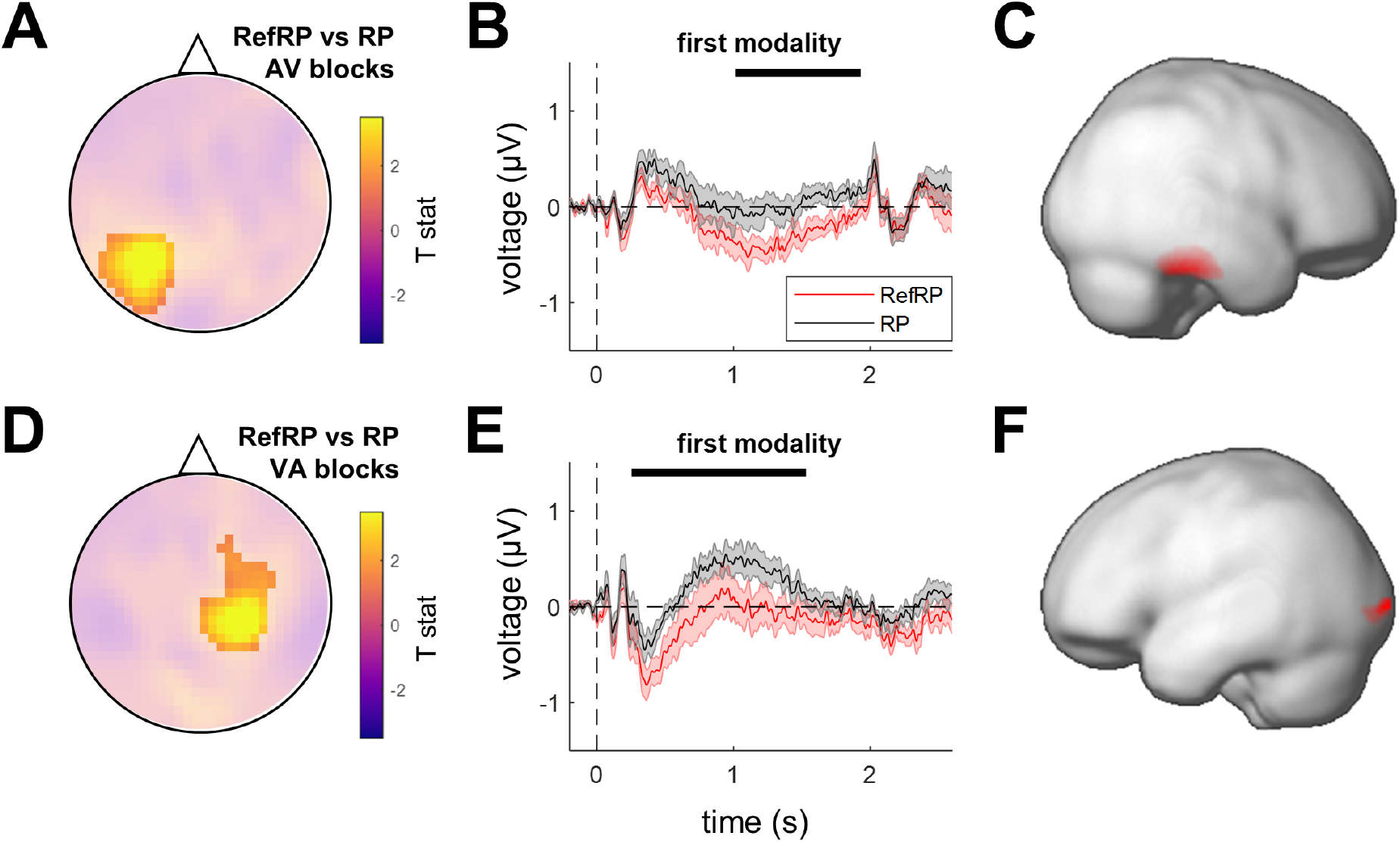
ERP effects of stimulus repetition. **(A)** Scalp topographies of repetition effects across trials (RefRP vs. RP stimuli), in the first modality of AV blocks. Highlighted significant clusters (p_FWE_ < 0.05). **(B)** Time courses of the effects of repetition across trials, in the first modality of AV blocks. Red: RefRP stimuli; black: RP stimuli. Shaded areas denote SEM across participants. Black bars denote the significant effect time windows. **(C)** Source estimates of the significant clusters in the AV blocks: right middle/inferior temporal gyrus (peak MNI [54 −24 −22]). **(D)** Scalp topographies of repetition effects across trials (RefRP vs. RP stimuli), in the first modality of VA blocks. Highlights as in (A). **(E)** Time courses of the effects of repetition across trials, in the first modality of VA blocks. Legend as in (B). **(F)** Source estimates of the significant clusters in the VA blocks: left inferior occipital gyrus (peak MNI [−22 −98 0]).

**Figure 4.**
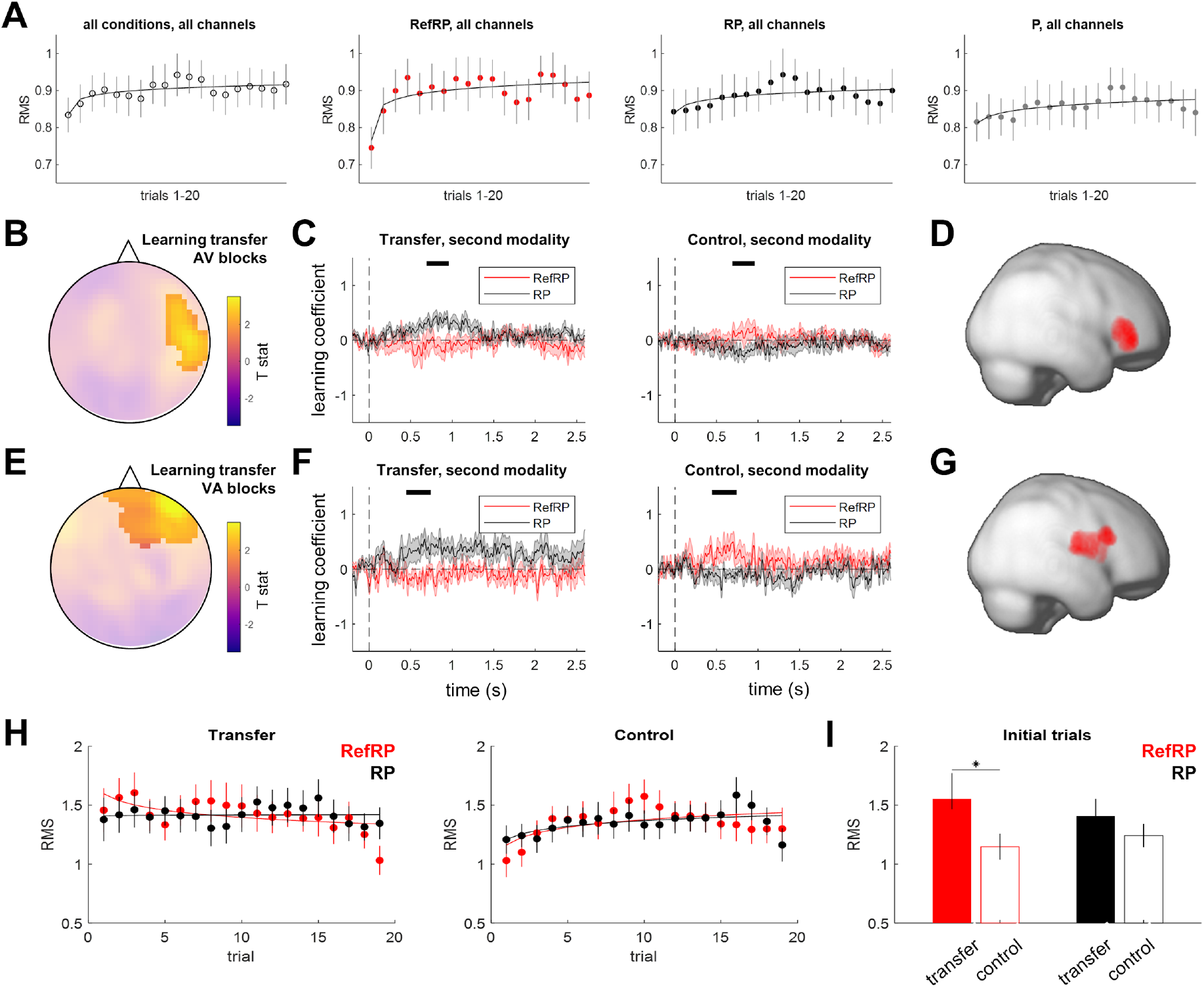
EEG learning curves reflect cross-modal transfer. **(A)** Learning curve estimation by logarithmic fitting. Plots show RMS of ERP responses (relative to baseline), averaged across channels and participants for all conditions (left), and for RefRP, RP, and P stimuli separately (middle/right). For display purposes, data were smoothed by a moving average of 3 points. Error bars denote SEM across participants. **(B)** Scalp topography of the cross-modal transfer effect (interaction: [RefRP vs. RP] x [Transfer vs. Control]) on EEG learning curves, in the second modality of AV blocks. Highlighted significant cluster (p_FWE_ < 0.05). **(C)** Time course of AV cross-modal transfer effect. Left: transfer condition, right: control condition. Red: RefRP stimuli; black: RP stimuli. Shaded areas denote SEM across participants. Black bars denote the significant interaction effect time windows. **(D)** Source estimates of the significant cluster: right inferior frontal gyrus (peak MNI [46 40 −4]). **(E)** Scalp topography of the cross-modal transfer effect in the second modality of VA blocks. Highlighted significant cluster (p_FWE_ < 0.05). **(F)** Time course of VA cross-modal transfer effect. Legend as in (B). **(G)** Source estimates of the significant cluster: right inferior frontal gyrus (peak MNI [52 14 20]). **(H)** Single-trial EEG amplitude (RMS). Left: transfer condition, right: control condition. Red: RefRP stimuli; black: RP stimuli. Solid curves denote logarithmic fits to the group average. Error bars denote SEM across participants. **(I)** Post-hoc tests of the EEG amplitude (RMS) averaged across the first three trials. Only the difference between RefRP EEG amplitudes (RMS) in transfer vs. control conditions was significant (p < 0.05).

**Figure 5.**
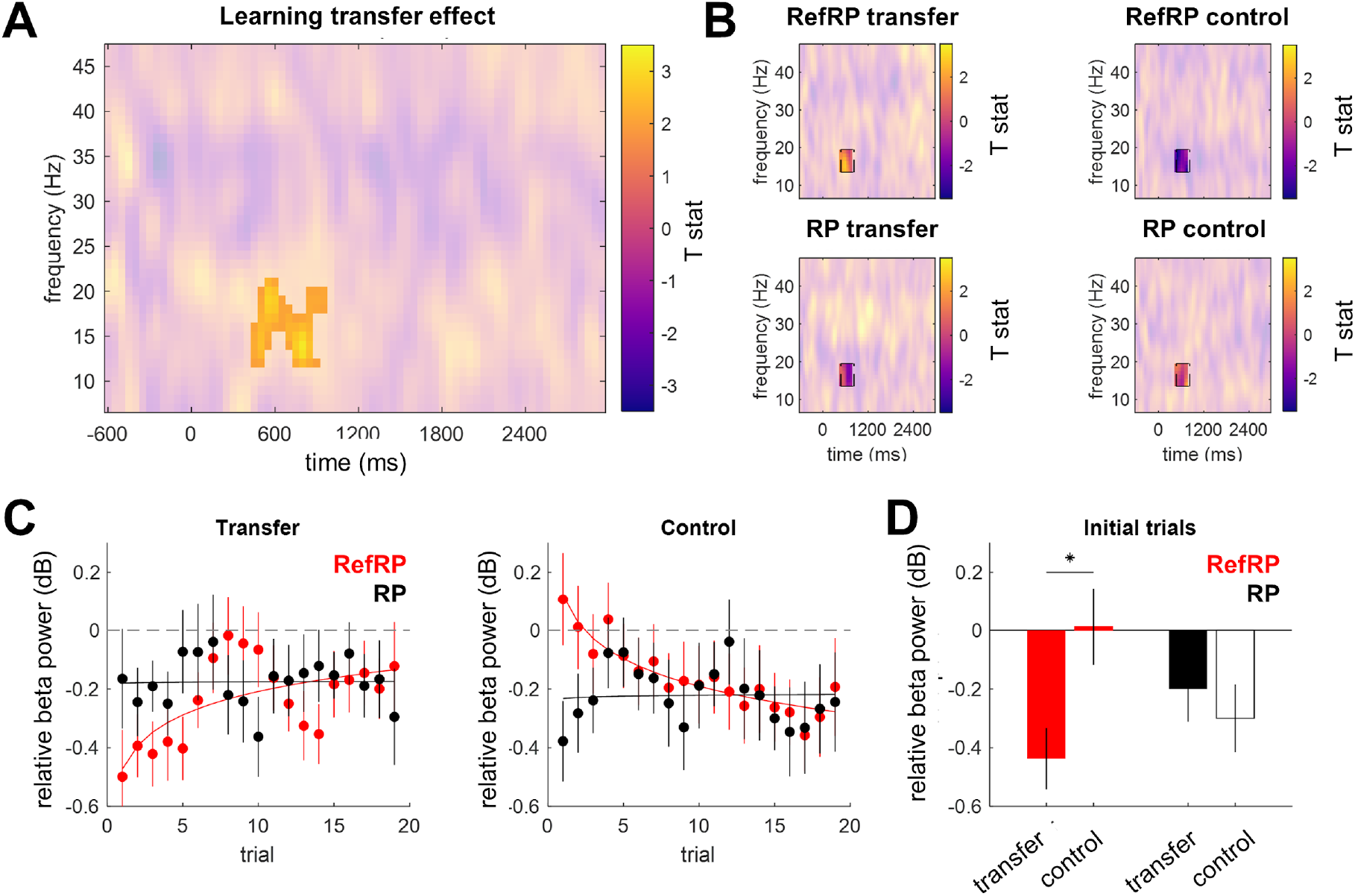
Frontal beta power reflects cross-modal transfer. **(A)** Time-frequency map of the cross-modal transfer effect (interaction: [RefRP vs. RP] x [Transfer vs. Control]) on single-trial changes (logarithmic coefficients) in rIFG activity, in the second modality of both AV and VA blocks. Highlighted significant cluster (*p_FWE_* < 0.05). **(B)** Time-frequency maps of each condition. Highlighted cluster denotes the time and frequency window (corresponding to the maximum range of the cluster depicted in Fig. 5A) used to obtain single-trial estimates in (C,D). **(C)** Single-trial frontal beta power. Left: transfer condition, right: control condition. Red: RefRP stimuli; black: RP stimuli. Solid curves denote logarithmic fits to the group average. Error bars denote SEM across participants. **(D)** Post-hoc tests of the frontal beta power, averaged across the first three trials. Only the difference between RefRP frontal beta power in transfer vs. control conditions was significant (*p* < 0.05).

## 3. Results

### 3.1 Behavioral effects

First, a mixed-design ANOVA on *d*’ confirmed a significant effect of stimulus type (*F*_1,22_ = 24.23, *p* < 0.001), modality (*F*_1,22_ = 1541.1, *p* < 0.001), and condition (*F*_1,22_ = 27.97, *p* < 0.001). A significant interaction for the stimulus type, modality, condition, and modality order was also observed (*F*_1,22_ = 4.556, *p* < 0.05). As the significant interaction was observed when the modality order was included, we ran separate repeated-measures ANOVA on each AV and VA group. First, overall performance in the auditory modality was better (that is, *d*’ values were higher) than in the visual modality for both AV and VA trial types (AV: *F*_1,11_ = 1140.37, *p* < 0.001; VA: *F*_1,11_ = 420.15, *p* < 0.001). Better performance for RefRP compared to RP was confirmed only in the VA group (*F*_1,11_ = 32.44, *p* < 0.001), and that of the AV group showed borderline significance (*F*_1,11_ = 4.54, *p* = 0.057). Due to the significant performance difference observed between A and V, further post-hoc ANOVAs were carried out on each modality separately. In the AV group, ANOVA on the first modality (Audition) confirmed that there was a significant performance difference between RP and RefRP (*F*_1,11_ = 6.53, *p* = 0.027) and no overall performance difference between Transfer and Control conditions (*F*_1,11_ = 0.33, *p* = 0.58). In contrast, an ANOVA on the second modality (Vision) revealed no significant performance difference between stimulus type (*F*_1,11_ = 0.20, *p* = 0.66) but a significant performance difference between condition (*F*_1,11_ = 28.28, *p* < 0.001) and a weak interaction between the stimulus type and condition (*F*_1,11_ = 4.33, *p* = 0.06). For the VA group, a post-hoc ANOVA on each modality confirmed that both modalities showed significantly greater performance for RefRP than RP (V: *F*_1,11_ = 14.05, *p* = 0.004; A: *F*_1,11_ = 29.86, *p* < 0.001), and for Transfer than Control (V: *F*_1,11_ = 182.18, *p* < 0.001; A: *F*_1,11_ = 180.18, *p* < 0.001). No effect of the condition or interaction between condition and stimulus type were observed (Fig. 2A), possibly due to the ceiling effect for task performance in the second modality (A). Overall, significant performance differences between RP and RefRP were observed in all cases except for the second modality of the AV group, which is the only case where a significant performance difference between Transfer and Control conditions was observed.

To investigate whether observed performance differences between stimulus type and modality type were due to a performance improvement of RefRP, especially for the Transfer condition, we computed a hit rate difference between RefRP and RP for each trial, for each modality order group. Unlike previous studies (e.g., Agus et al., 2010; Kang et al., 2018), we did not observe a rapid performance improvement (within 5-10 trials) in all cases, possibly due to a limited number of test blocks (see Discussion). However, participants still showed greater performance difference in the second modality of the Transfer condition vs. Control condition, at least for the AV group (*t*(11) = −2.89, *p* < 0.05, Bonferroni correction; Fig. 2B), while no significant performance difference was observed in all other cases, i.e., the first modality in both groups as well as the second modality in the VA group (*p* > 0.05), which further supports the hypothesis that behavioural effects of pattern learning transfer should be most robust from audition to vision.

### 3.2 Stimulus repetition modulates ERP amplitude

In the analysis of the effects of stimulus repetition across trials (RefRP vs. RP) on ERP amplitude, two significant spatiotemporal clusters were identified (Fig. 3AD). In the first modality of the AV blocks (i.e., for auditory stimuli), RefRP stimuli differed in amplitude from RP stimuli over posterior channels, during the second stimulus segment (latency range 1110-1900 ms, cluster-level p_FWE_ < 0.001, F_max_ = 34.27, Z_max_ = 5.37), while for the first modality of the VA blocks (i.e., for visual stimuli), RefRP stimuli differed in amplitude from RP stimuli over central channels during both stimulus segments (latency range 372-1442 ms, cluster-level p_FWE_ < 0.001, F_max_ = 29.21, Z_max_ = 4.99). The remaining effects did not yield significant clusters. In the source reconstruction of the two significant clusters (Fig. 3CF), the difference in auditory RefRP and RP stimuli was attributed to the right middle/inferior temporal gyrus (MNI peak: [54 −24 −22], cluster-level p_FWE_ = 0.047, small-volume corrected, T_max_ = 2.42, Z_max_ = 2.37), in the immediate vicinity of secondary auditory regions, while the difference in visual RefRP and RP stimuli was attributed to the left inferior occipital gyrus (MNI peak: [−22 −98 0], cluster-level p_FWE_ = 0.047, small-volume corrected, T_max_ = 2.30, Z_max_ = 2.26), in the early visual cortex.

### 3.3 Cross-modal transfer modulates EEG learning curves

In the analysis of the effects of cross-modal transfer (interaction: [RefRP vs. RP] x [Transfer vs. Control]) on EEG learning curves, two significant clusters were identified. In the second modality of the AV blocks (i.e., for visual stimuli), there was a significant interaction between stimulus (RefRP vs. RP) and condition (Transfer vs. Control) over right-lateralized central electrodes, during the first stimulus segment (latency range 704-942 ms, cluster-level *p_FWE_* = 0.042, *F_max_* = 15.41, *Z_max_* = 3.63; Fig. 4BC), while for the second modality of the VA blocks (i.e., for auditory stimuli), the same interaction yielded a significant cluster over right-lateralized frontal electrodes, also during the first stimulus segment (latency range 476-700 ms, cluster-level *p_FWE_* = 0.003, *F_max_* = 23.30, *Z_max_* = 4.47; Fig. 4EF). In the source reconstruction of these clusters, both significant interaction effects were attributed to source activity in the right inferior frontal gyrus (rIFG; AV blocks, MNI peak: [46 40 −4], cluster-level *p_FWE_* = 0.049, small-volume corrected across voxels, *T_max_* = 3.17, *Z_max_* = 3.05; Fig. 4D; VA blocks, MNI peak: [52 14 20], cluster-level p_FWE_ = 0.049, small-volume corrected across voxels, *T_max_* = 3.67, *Z_max_* = 3.50; Fig. 4G).

A further inspection of the interaction effect (Fig. 4HI) revealed that, for the initial three trials of the second modality, it was driven by the significant difference in ERP amplitude between RefRP stimuli presented in Transfer vs. Control conditions (*t_23_* = 2.38, *p* = 0.026), with no significant differences between the remaining stimulus/condition pairs (all other *p* > 0.2).

### 3.4 Cross-modal transfer modulates frontal beta power

For the analysis of time-frequency activity, we estimated source activity localized to the rIFG region, and tested whether any time-frequency clusters are sensitive to cross-modal transfer (interaction effect) as identified above for ERPs. This analysis revealed that, across AV and VA blocks, beta-band power was modulated by cross-modal transfer during the first stimulus segment (latency range 500-800 ms, frequency range 15-20 Hz, cluster-level *p_FWE_* = 0.016, *T_max_* = 3.25, *Z_max_* = 3.19; Fig. 5AB). The main effect of block order (AV vs. VA) was not significant. A closer inspection of this interaction effect for the first three trials (Fig. 5CD) revealed that, as in the case of ERPs, it was driven by the significant difference beta power between RefRP stimuli presented in Transfer vs. Control conditions (*t_23_* = −2.59, *p* = 0.016), with no significant differences between the remaining stimulus/condition pairs (all other *p* > 0.1).

### 3.5 Neural signatures of learning transfer contribute to behaviour

Across participants, the behavioural benefits in sensitivity to RefRP vs. RP stimuli were explained in an ANCOVA by a combination of EEG signatures of learning transfer (difference in logarithmic fit coefficients for RefRP vs. RP stimuli based on EEG amplitudes; see Fig. 4BC and Fig. 4EF), beta-band signatures of learning transfer (difference in logarithmic coefficients for RefRP vs. RP stimuli based on beta power; Fig. 5B), experimental condition (Transfer vs. Control), and modality order (AV vs. VA). Here, for the behavioural benefit measure, we used raw *d*’ values (before rescaling; see Section XYZ) to preserve the individual differences. Specifically, there was a significant three-way interaction between the continuous covariate representing the EEG signatures of learning transfer, and the categorical factors of condition and order (*F_1,31_* = 4.36; *p* = 0.045), as well as a significant four-way interaction between the continuous covariates representing the EEG and beta-band activity signatures of learning transfer, and the categorical factors of condition and order (*F_1,31_* = 5.02, *p* = 0.032). The remaining main and interaction effects were not significant (all *p* > 0.1). A further inspection of the significant interaction effects revealed that, in the Transfer condition of the AV group, behavioural sensitivity benefits were subject to a significant main effect of EEG signatures of learning transfer (*F_1,8_* = 5.52, *p* = 0.047, *β* = 0.425), and a significant interaction effect between EEG and beta-band signatures of learning transfer (*F_1,8_* = 7.67, *p* = 0.024, *β* = −0.262). These effects were not significant in the Control condition or in the VA group (all *p* > 0.05).

## 4. Discussion

In this study we show that the temporal patterns of occasionally repeated stimuli (RefRP) are learned and exploited, both when these stimuli are presented within a particular modality (in both audition and vision), as well as when the temporal patterns are learned in one modality and presented in another. First, participants had better behavioural performance (within-trial repetition detection) for RefRP stimuli, presented several times during the experiment, than for RP stimuli, only presented once. These performance benefits were observed in both audition and vision as the first modality, consistent with previous studies using the same paradigm (Kang:2018ee; Kang et al., 2017; Gold et al., 2014; Bale et al., 2017), and regarded as an index of learning for RefRP. A qualitatively similar performance improvement in both modalities was achieved by controlling presented temporal patterns at a low pulse rate with a fixed minimum interval (Rammsayer et al., 2015; Rammsayer and Pichelmann, 2018).

The ERP difference observed between RefRP and RP in the first modality further confirmed that stimulus repetition across trials modulates the processing of that stimulus. We observed a significant ERP difference between RefRP and RP in both audition and vision, complementing previous studies using similar paradigms which reported a modulation of neural responses to stimulus repetition in audition (Luo et al., 2013; Andrillon et al., 2015). However, while previous studies focused on comparing RefRP stimuli to completely random sequences (the equivalent of P stimuli in our study) and did not report significant differences between RefRP and RP stimuli on ERP responses, in this study we focused *a priori* on comparing RefRP and RP stimuli, since there are fewer confounding factors such as motor response preparation. Furthermore, while previous studies used Gaussian noise as stimuli, in the present study we used random temporal patterns of discrete brief stimuli. Our rationale for using relatively slow (average pulse rate 7 Hz) patterns of discrete stimuli was to minimize the absolute differences in temporal acuity between audition and vision and ensure that each pulse gap could be clearly perceived by both modalities (Kang et al., 2018). However, as a result the sequence used in the present study was simpler and more salient than frozen noise stimuli, possibly augmenting differences in neural responses to RefRP and RP stimuli. A source localisation of these effects showed that, in audition, differences in ERPs evoked by RefRP vs. RP stimuli were primarily due to activity in or near the auditory cortex while, in vision, these differences originated from the visual cortex. This is contradictory to some of the previous studies suggesting an involvement of auditory regions in temporal processing of both auditory and visual stimuli (Guttman et al., 2005; Kanai et al., 2011), or an involvement of visual regions in mnemonic processing of both auditory and visual stimuli (Wolff et al., 2020). Instead, our findings of modality-specific neural dynamics modulated by repeated presentation of the same temporal pattern are more consistent with the intrinsic model, according to which temporal information is coded in modality-specific sensory regions (Ivry et al., 1988; Ivry and Schlerf, 2008; Buonomano and Maass, 2009).

Crucially, beyond the effects of occasional pattern repetition on stimulus processing within a given modality, we also observed behavioural and neural correlates of learning transfer across modalities. In the behavioural data, there was a trend for better performance achieved for RefRP stimuli which were first presented in audition and whose temporal characteristics were transferred to vision. While no such finding was observed for transfer from vision to audition, this could be due to a behavioural ceiling effect in audition. Also, a limited number of test blocks run for the EEG recording could have caused the unbalanced task performance between conditions for the first modality, vision, with greater variability. However, since such unbalanced task performance in vision was observed only for early trials and disappeared for later trials, it would not have affected our investigation of the transfer effect. Transfer effects of learning across modalities have not been consistent across previous studies. Some studies reported that training in audition improves visual temporal processing but not vice versa (Bratzke et al., 2012; Barakat et al., 2015), while other studies showed a generalization of temporal perceptual learning from vision to audition (Bueti and Buonomano, 2014) or even no transfer effect at all (Lapid et al., 2008). It should be noted, however, that because we recorded neural responses concurrently, the test blocks were longer and as a result a smaller number of behavioural blocks were presented compared to the previous studies, possibly decreasing the behavioural sensitivity to learning transfer.

Nevertheless, the analysis of neural responses showed robust correlates of learning transfer across modalities, in both directions (from audition to vision and from vision to audition). Previous auditory imaging studies in humans under the same paradigm focused on the inter-trial coherence as an index of neural activity modulated by the re-occurrence of reference sequences (e.g., Luo et al., 2013; Andrillon et al., 2015). However, the inter-trial coherence by definition averages information across trials. In the present study, since we were primarily interested in learning transfer, we estimated a neural learning curve to repeated presentations of RefRP stimuli across trials, based on a logarithmic fit of trial-by-trial responses. Our choice of logarithmic fitting, as opposed to other (e.g. linear and exponential) fits analysed here, is consistent with previous work on statistical learning in adults (Siegelman et al., 2018) and infants (Choi et al., 2020) which has shown that logarithmic fits efficiently describe learning curves. This allowed us to quantify the gradual changes in neural responses, tapping more directly into the learning mechanisms. Using these methods, we found that learning transfer of RefRP temporal patterns from one modality to another, for both AV and VA groups, were reflected by a significant difference in the ERP learning curves between RefRP stimuli presented in the second modality in the transfer condition compared to the control condition, whereas no such difference was observed for RP stimuli. A significant interaction effect of stimulus (RefRP vs. RP) and condition (transfer vs. control) on ERP learning curves was observed in the second modality for both AV and VA groups. Importantly, these effects were driven by the initial few trials - and, unlike the first modality, in which we observed overall differences between ERP amplitudes evoked by RefRP and RP stimuli, here the average differences in ERP amplitudes were not significant. Furthermore, in the AV group, in which we observed significant behavioural benefits of learning transfer, these benefits were explained by the ERP signatures of learning transfer (i.e., EEG-based learning curves predicted the size of the behavioural benefit in the Transfer condition, but not in the Control condition). This indicates that there is a distinct difference in neural responses driven by memory transfer across modalities, immediately after presenting a previously learned stimulus in a new modality, and largely limited to the initial stimulus presentations.

Interestingly, for both AV and VA groups, source localization revealed that the learning transfer effect was associated with activity modulations in the right IFG, typically associated with working memory, attention, and detection of relevant targets, regardless of modality (Linden et al., 1999; Corbetta and Shulman, 2002; Hampshire et al., 2010). Furthermore, within the rIFG cluster, significant signal modulation due to the learning transfer effect was specifically observed in the beta-band, also implicated in working memory function (Spitzer and Blankenburg, 2012; Lautz et al., 2017; Gelastopoulos et al., 2019). Also, the interaction of beta-band and ERP-based learning curves could be related to the behavioural benefits of learning transfer, suggesting that the interplay between both kinds of neural signatures has behavioural relevance. Previous work has identified the right prefrontal beta-band activity as a neural correlate of working memory maintenance independent of modality or specific stimulus type (Spitzer et al., 2014; Wimmer et al., 2016), and mounting evidence suggests that prefrontal beta is linked to reactivating working memory contents (Spitzer and Haegens, 2017). In this context, our findings are consistent with the notion that learning transfer requires a reactivation of a previously learned temporal pattern in working memory, possibly matching the temporal pattern in a new sensory format to the pattern in the previously-learned modality.

## 5. Conclusion

The present study suggests that both modality-specific and modality-general mechanisms mediate temporal information processing, depending on whether temporal sequences are processed in one modality or transferred across modalities. Different effects of the reference sequence reoccurrence on neural activity during the first and second modality learning phases suggest dissociable information processing mechanisms between the two phases. A dominant involvement of modality-specific cortical regions during the first phase could be explained by the intrinsic model, according to which temporal information is initially coded in the contextually relevant regions (Buonomano and Maass, 2009). However, higher-order processing of information, i.e. transferring the temporal pattern learned in one modality to another modality, relies on reactivating the relevant stimulus contents in working memory (Spitzer and Haegens, 2017). Similar neural effects of learning and learning transfer independent of modality (audition, vision) and order (audition-to-vision and *vice versa*) suggests that general mechanisms might be at play when learning temporal patterns in different sensory modalities.

## Acknowledgements

This work has been supported by the European Commission’s Marie Skłodowska-Curie Global Fellowship (750459 to R.A.), the Hong Kong General Research Fund (11100518 to R.A. and J.S.), a grant from European Community/Hong Kong Research Grants Council Joint Research Scheme (9051402 to R.A. and J.S.), and Fyssen Foundation (post-doctoral study grant to H.K.). We would like to thank Reuben Chaudhuri for help with data acquisition, and Sir Colin Blakemore for helpful discussions.

## Author contributions

Conceptualization and Methodology: HK, RA, CHC, VGR, and JWHS; Data collection: HK, RA, CHC, DC, and JWHS; Data analyses and Visualisation: HK and RA; Writing original draft: HK and RA; Reviewing and editing: HK, RA, DC, VGR, and JWHS; Supervision: JWHS.

